# AutoRNAseq: Automated Bulk RNA-seq Analysis Pipeline

**DOI:** 10.64898/2026.04.21.719844

**Authors:** Josh Loecker, Brandt Bessell, Bhanwar Lal Puniya, Tomáš Helikar

## Abstract

**Summary:** Improved accessibility of high-throughput RNA sequencing has increased the amount of data generated each year. This increase in data creates a need for reproducible pipelines that can process RNA-seq data consistently across experiments. AutoRNAseq addresses this need by providing a Snakemake-based workflow for bulk RNA-seq analysis by automating data retrieval, quality control, and gene quantification. Unlike existing RNA-seq workflows that require users to coordinate multiple pipelines and pre-configure reference data, AutoRNAseq provides a single, end-to-end workflow that automates data acquisition, reference preparation, quality control, alignment, and quantification with minimal user intervention. AutoRNAseq is applicable to any domain requiring consistently processed RNA-seq datasets, including bioinformatics, computational biology, and drug-response studies.

**Availability and Implementation:** AutoRNAseq is implemented in Snakemake and available at https://gitlab.com/unebraska/lagbh-public/autornaseq. Documentation and example configuration files are provided in the GitLab README file and this paper’s Supplementary Information. The code to reproduce the statistics presented here is in the GitLab repository under the “publication” folder.

## Introduction

Genomics has experienced a significant expansion in sequencing capacity over the past decade.^1^ Advances in next-generation sequencing technology have dramatically reduced per-sample costs, enabling researchers to generate increasingly large and complex RNA-seq datasets. This ease of sequencing has accelerated biological discovery but has also created a “reproducibility crisis”, where subtle differences in software versions or parameters can substantially affect results.^2^ Workflow managers, such as Snakemake^3^, address this by combining Python’s flexibility with file-based dependency tracking, allowing explicit declaration of software dependencies through containers or Conda environments.

Despite these tools, many RNA-seq pipelines require substantial infrastructure setup and manual preparation. For example, the `nf-core/rnaseq` pipeline requires users to provide pre-formatted genome FASTA and GTF files, which can introduce errors if generated incorrectly.^4,5^ Here, we present AutoRNAseq, a Snakemake-based bulk RNA-seq pipeline that significantly minimizes user intervention while maximizing reproducibility. AutoRNAseq directly fetches data from NCBI GEO accessions or reads from local repositories, prepares reference genomes from Ensembl, and produces gene count matrices for downstream analysis, such as differential expression analysis.

### Implementation Details

AutoRNAseq is organized into the following five stages:

1. Workflow setup and reference genome preparation: AutoRNAseq parses the provided configuration file (discussed in the Pipeline Behavior section below) and writes metadata files describing sample layout and library preparation. Additionally, the reference FASTA and annotation (GTF) files for the specified release will be downloaded, and their related indices will be built using STAR. Following this, the corresponding transcriptome FASTA file is generated.
2. Data downloading: if the user chooses to use publicly available data, the `prefetch` command-line tool will be used to download SRR accession information from NCBI’s Gene Expression Omnibus (GEO).
3. Data preprocessing and quality control: AutoRNAseq will, optionally, perform read trimming with Trim Galore!.^6^ Quality control processing with FastQC is performed on the raw and, if available, trimmed, FASTQ files.^7^ Finally, this section of the pipeline will scan for possible contamination using the FastQ Screen command-line tool against the following genomes: adapter sequences, Arabidopsis, Drosophila, E. coli, human, lambda phages, mitochondria, mouse, PhiX phages, rat, vectors, worm, yeast, and rRNA.^8^
4. Alignment and quantification: Reads are aligned to the genome using STAR, producing a coordinate-sorted BAM file for downstream processing. Salmon then estimates transcript- and gene-level abundances.^9,10^ Although lightweight transcriptome-mapping tools (e.g., Salmon and Kallisto) offer faster quantification, STAR was selected because it supports alignment-level quality control, visualization, and optional downstream analysis.
5. Metric calculation and report generation: AutoRNAseq computes RNA-seq metrics and insert sizes using Picard. A comprehensive quality control report is generated with MultiQC.^11,12^ Additionally, fragment sizes, insert sizes, RNA-seq metrics, and gene count quantifications are copied into a directory structure directly readable by COMO, a Jupyter Notebook-based tool for automated constraint-based metabolic modeling, discussed later.^13^

### Pipeline Behavior

AutoRNAseq provides two methods for data input, supporting both public datasets and local sequencing files. Both approaches require a metadata file containing sample information, including columns for GEO SRR accession numbers, sample names, end type (single-or paired-end), and sequencing method (total RNA or polyA). First, if SRR accession numbers are used, the pipeline will automatically download and process each dataset.^14^ This approach helps combine multiple studies for additional downstream analysis. The second input method processes local FASTQ files. This approach suits individuals who have their own set of sequencing data to analyze. In this method, the first column of the metadata file should be left blank, and the directory path to the FASTQ files should be specified in the configuration file under the `LOCAL_FASTQ_FILES` key. The FASTQ filenames should be prefixed with the respective “sample” identifier, enabling automatic association between files and sample metadata.

The pipeline needs a configuration file that controls its behavior. Key parameters include the path to the metadata CSV file and output directories. More granular control is also provided over which processing steps are executed: adapter trimming, contaminant screening, RNA-seq metrics calculation, insert size collection, and fragment size collection. The reference genome configuration requires specifying the taxonomy ID and Ensembl release version, which allows AutoRNAseq to automatically download the required reference genome and pre-computed alignment indices from Ensembl.^15^ The outputs of this include the genome FASTA, annotation GTF, STAR’s genome index, FASTA transcriptome (cDNA), and the Salmon index. Lastly, benchmarks are automatically calculated to track execution time, CPU usage, and memory usage for each rule.

Once users configure the above, executing the pipeline is straightforward. Two profiles are provided by default, for local or SLURM workflow execution. For local execution, use the command `snakemake --profile profiles/local`. Alternatively, if running the workflow on a computing cluster, use the command `snakemake --profile profiles/cluster`. This will automatically populate Snakemake with the required job names, account details, and resources to submit jobs to the cluster. Jobs are submitted to the cluster based on its available resources. Because of this, Snakemake does not submit a single job to the cluster requesting the sum of each rule’s runtime, memory, and CPU requirements, as this would unnecessarily inflate resource usage. Instead, Snakemake submits a single job per rule, using the resource limits we have carefully tweaked to increase the job submission rate and decrease the total runtime.

The figure below shows the information flow within AutoRNAseq. At a minimum, at least one red input must be provided (local FASTQ files or SRR accession values). After this, any blue step can be skipped by setting its corresponding configuration value to False.

### Comparison to Other Tools

RNA-seq pipelines differ in computational robustness, supporting downstream biological interpretation and integration with multi-omic modeling workflows. The most popular and comprehensive RNA-seq analysis pipeline is `nf-core/rnaseq`, which provides extensive tool integration and rigorous testing.^4^ However, `nf-core/rnaseq` relies on a separate workflow (`nf-core/fetchngs`) for SRA downloads, requiring users to coordinate two distinct pipelines and separate terminal commands for complete data processing.^5^ In contrast, AutoRNAseq integrates data acquisition and alignment in a single workflow, streamlining the process for individuals working primarily with GEO/SRA data.

Most RNA-seq pipelines are optimized for differential gene expression analysis and reporting. In contrast, AutoRNAseq was explicitly designed to serve as a bridge between transcriptomic preprocessing and downstream systems-level modeling workflows, such as genome-scale metabolic modeling, rather than as a standalone differential expression pipeline. Namely, AutoRNAseq outputs results in a format that is compatible with the “Constraint-based Optimization of Metabolic Objectives” (COMO) tool, a Jupyter notebook-based pipeline for multi-omic data integration in metabolic modeling and drug discovery.^13^ COMO requires processed gene expression matrices as input, which AutoRNAseq generates automatically in its alignment and quantification steps. This integration allows individuals to transition quickly and easily from bulk RNA-seq analysis to reconstructing context-specific, constraint-based metabolic models. The combination of these tools allows for multi-scale studies that connect transcriptomics, metabolic predictions, and drug target identification.

## Results & Use Case

To demonstrate AutoRNAseq capabilities, we applied the pipeline to a use case involving multiple human RNA-seq studies from the Sequencing Quality Control Consortium.^16^ This analysis evaluated the pipeline’s ability to handle diverse experimental designs, but focused primarily on paired-end, total RNA preparation.

The analysis incorporated thirty-two SRR IDs from the GSE47792 identifier (Supplementary Table S1). The samples in this table encode biological context, allowing multiple experiments to be aligned simultaneously. The data are encoded as follows: cell type or tissue, study identifier, and run number. If studies include technical replicates, a lowercase ‘r’ followed by the replicate number should be appended to the sample name. This structured naming facilitates downstream analysis while maintaining clear file tracking.

The pipeline was executed on a workstation with 24 cores and 256 GB RAM using the command `snakemake --profile profiles/cluster`. We validated pipeline accuracy by comparing gene counts generated by AutoRNAseq with those from the `nf-core/rnaseq` pipeline using the same Ensembl reference genome, release 115. Figure 1B shows the results of these statistics and Supplementary Table S2 presents the values for graph creation. We compared log-transformed per-gene counts between AutoRNAseq and `nf-core/rnaseq`. For each sample, all Pearson correlation coefficients exceed 0.9839 (P-values < 0.05), indicating a strong linear relationship between the two pipelines. Similarly, Spearman correlations are above 0.8770 (P-values < 0.05), indicating that genes are ranked similarly between the pipelines. Finally, the Kolmogorov-Smirnov statistics are all approximately 0.0025 (P-values > 0.05), suggesting that we cannot reject the null hypothesis that gene count distributions from each pipeline are the same. Thus, taken together, AutoRNAseq produces results similar to those of the `nf-core/rnaseq` pipeline.

**Figure 1:**
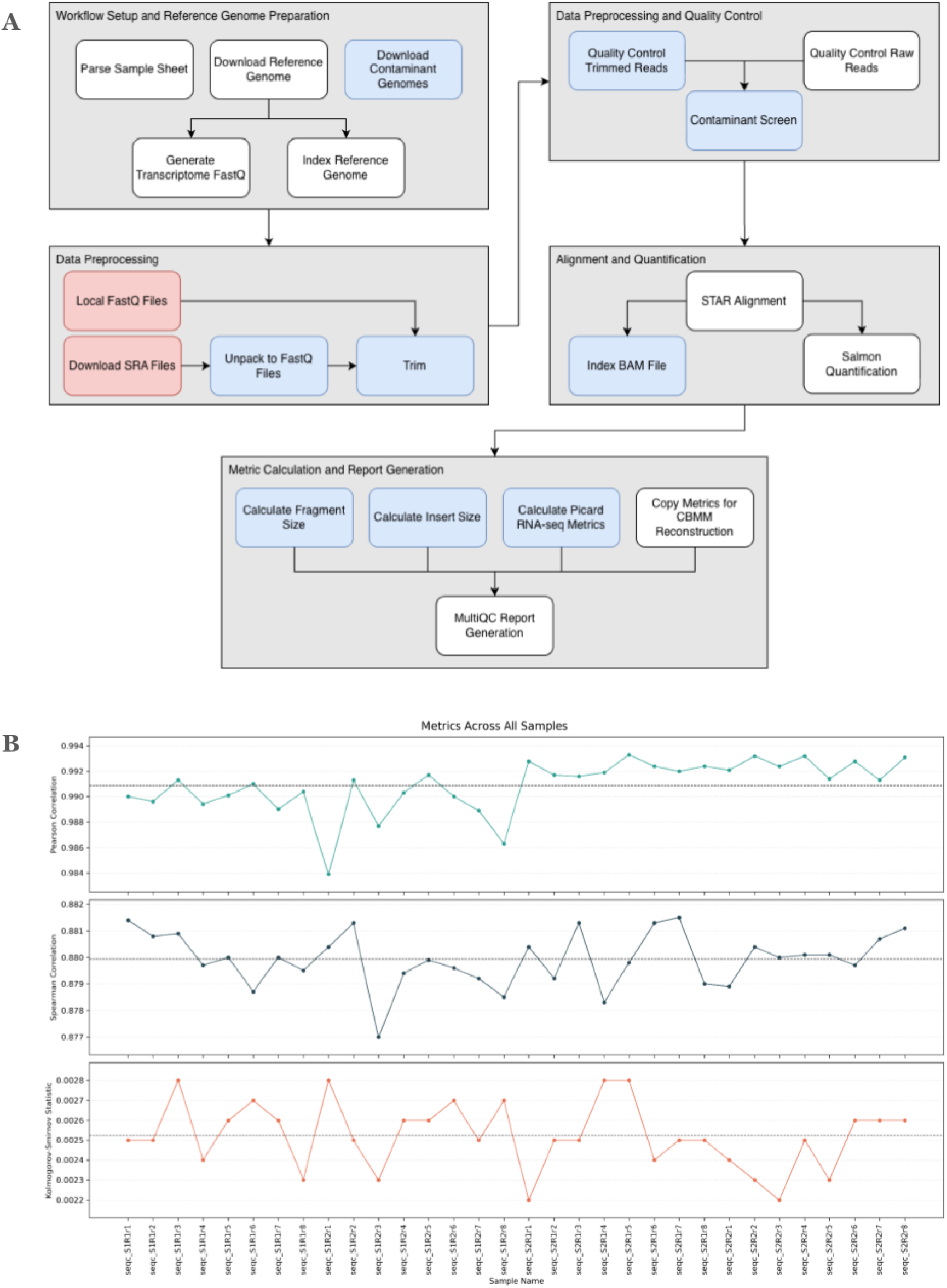
Information flow and statistical comparisons. Panel 1A): AutoRNAseq’s information flow. At least one red input is required (local FASTQ files or NCBI GEO SRR accession numbers), and any blue processing step can be skipped by modifying the related setting. Panel 1B): Pearson, Spearman, and Kolmogorov-Smirnov statistics between AutoRNAseq and nf-core/rnaseq.

## Conclusion

AutoRNAseq provides a reproducible, accessible solution for bulk RNA-seq alignment that addresses key challenges in modern genomics research. By integrating data acquisition, quality control, alignment, and quantification in a single Snakemake workflow, the pipeline reduces technical barriers while maintaining rigorous reproducibility standards. Dual input modes accommodate both public data meta-analyses and local sequencing projects, while Conda-based dependency management ensures consistent results across computing environments. Our validation results demonstrate that the pipeline outputs closely align with those of the existing `nf-core/rnaseq` tool. AutoRNAseq provides an easy-to-use, reproducible RNA-seq alignment pipeline that requires minimal configuration. Documentation, example configuration files, and the complete source code are available at the GitLab repository.

