## Supplementary Information for "AutoRNAseq: Automated Bulk RNA-seq Analysis Pipeline"

### Supplementary Material: AutoRNAseq – Automated Metabolic Model Preparation Pipeline

#### Configuration and Workflow Details

##### Environment Variables

By default, AutoRNAseq submits SLURM cluster jobs to the first account associated with the logged-in user, as shown in the GitLab repository in the file `profiles/cluster/submit.sh`. If users would like to manually define the SLURM account, the environment variable `SNAKEMAKE_SLURM_ACCOUNT` can be set before starting the pipeline. For example, the variable can be set in a single line as follows (by changing `my-account-name` to the SLURM account):

```
SNAKEMAKE_SLURM_ACCOUNT=my-account-name snakemake --profile profiles/cluster
```

Or, if users would prefer to set the variable and then execute the pipeline in two separate commands:

```
export SNAKEMAKE_SLURM_ACCOUNT=my-account-name

snakemake --profile profiles/cluster
```

##### Metadata CSV Format

AutoRNAseq requires a CSV metadata file with four columns: sample name, SRR ID, end type, and preparation method. The file must include a header row. Below is the complete metadata used in our validation analysis.

Table S1) Complete Metadata CSV for Validation Study

Note: The table header should only be included once, but it may be shown multiple times in this file for ease of tracking header names with columns

This table can be converted to a CSV file and included in AutoRNAseq's `SAMPLE_FILEPATH` file to reproduce the work shown here.

| srr | sample | endtype | prep_method |
|---|---|---|---|
| SRR896663 | seqc_S1R1r1 | PE | total |
| SRR896665 | seqc_S1R1r2 | PE | total |
| SRR896667 | seqc_S1R1r3 | PE | total |

| srr | sample | endtype | prep_method |
|---|---|---|---|
| SRR896669 | seqc_S1R1r4 | PE | total |
| SRR896671 | seqc_S1R1r5 | PE | total |
| SRR896673 | seqc_S1R1r6 | PE | total |
| SRR896675 | seqc_S1R1r7 | PE | total |
| SRR896677 | seqc_S1R1r8 | PE | total |
| SRR896664 | seqc_S1R2r1 | PE | total |
| SRR896666 | seqc_S1R2r2 | PE | total |
| SRR896668 | seqc_S1R2r3 | PE | total |
| SRR896670 | seqc_S1R2r4 | PE | total |
| SRR896672 | seqc_S1R2r5 | PE | total |
| SRR896674 | seqc_S1R2r6 | PE | total |
| SRR896676 | seqc_S1R2r7 | PE | total |
| SRR896678 | seqc_S1R2r8 | PE | total |
| SRR896679 | seqc_S2R1r1 | PE | total |
| SRR896681 | seqc_S2R1r2 | PE | total |
| SRR896683 | seqc_S2R1r3 | PE | total |
| SRR896685 | seqc_S2R1r4 | PE | total |
| SRR896687 | seqc_S2R1r5 | PE | total |
| SRR896689 | seqc_S2R1r6 | PE | total |
| SRR896691 | seqc_S2R1r7 | PE | total |
| SRR896693 | seqc_S2R1r8 | PE | total |
| SRR896680 | seqc_S2R2r1 | PE | total |
| SRR896682 | seqc_S2R2r2 | PE | total |
| SRR896684 | seqc_S2R2r3 | PE | total |
| SRR896686 | seqc_S2R2r4 | PE | total |
| SRR896688 | seqc_S2R2r5 | PE | total |
| SRR896690 | seqc_S2R2r6 | PE | total |
| SRR896692 | seqc_S2R2r7 | PE | total |
| SRR896694 | seqc_S2R2r8 | PE | total |

**Column Descriptions:**

- sample: Unique identifier encoding cell type, study, and run/replicate information
- srr: NCBI SRA Run identifier for downloading from GEO
- endtype: "PE" for paired-end sequencing, "SE" for single-end sequencing
- prep_method: "mrna" for polyA-selected RNA, "total" for total RNA

Table S2) Pipeline Rules/Steps

Rules ending in `_paired` process paired-end reads, and rules ending in `_single` process single-end reads.

| Rule Name | Notes |
|---|---|
| `copy_config` | Copies the configuration file for future reference |
| `preroundup` | Prepares metadata for downstream rules |
| `download_genome` | Downloads the specified reference genome |
| `download_contaminant_genomes` | Downloads additional genomes for contamination quality control |
| `star_index_genome` | Indexes the reference genome |
| `generate_transcriptome_fasta` | Generates a transcriptome file for Salmon quantification |
| `prefetch` | Downloads SRA files from NCBI; only executed if not using local FASTQ files |
| `fastq_dump_paired`<br>`fastq_dump_single` | Converts SRA files to FASTQ files; only executed if not using local FASTQ files |
| `qc_raw_fastq_paired`<br>`qc_raw_fastq_single` | Performs `fastqc` for quality control on raw FASTQ files. |
| `trim_paired`<br>`trim_single` | Performs `trim-galore` on raw FASTQ files |
| `qc_trim_fastq_paired`<br>`qc_trim_fastq_single` | Performs `fastqc` for quality control on trimmed FASTQ files |
| `align` | Aligns the raw or trimmed FASTQ files to the reference genome |
| `index_bam_file` | Indexes the resulting aligned BAM file |
| `salmon_quantification` | Quantifies gene counts |
| `contaminant_screen_paired`<br>`contaminant_screen_single` | Checks the raw or trimmed FASTQ files for contamination using `fastq_screen` |
| `fragment_size` | Calculates fragment sizes for downstream analysis (e.g., FPKM calculations [not performed in AutoRNAseq]) |
| `insert_size` | Calculates insert sizes for downstream analysis (e.g., FPKM calculations [not performed in AutoRNAseq]) |

| Rule Name | Notes |
|---|---|
| `rnaseq_metrics` | Calculates metrics using `picard CollectRnaSeqMetrics` |
| `copy_fragment_size`<br>`copy_insert_size`<br>`copy_rnaseq_metrics`<br>`copy_gene_counts` | Copies the output from the related rule into a COMO-specific directory for downstream constraint-based metabolic model building (not performed in AutoRNAseq) |
| `multiqc` | Aggregates all quality-control output into a single HTML file for analysis |

### Validation Results

#### Gene Count Correlation Analysis

We compared gene counts generated by AutoRNAseq with those from the `nf-core/rnaseq` pipeline. Correlation was assessed using Pearson, Spearman, and Kolmogorov-Smirnov correlations. Statistics summaries (min, max, mean, median) are shown at the bottom of the table.

<u>Table S3) Validation Metrics by Sample</u>

Note: All statistics were performed on log1p-transformed gene count data. The p-values for Pearson and Spearman correlations were below the precision thresholds of NumPy's Float64 data type ($5e^{-324}$), and are reported as 0.

| Sample | Pearson | | Spearman | | Kolmorogorov-Smirnov | |
|---|---|---|---|---|---|---|
| | Statistic | P-value | Statistic | P-value | Statistic | P-value |
| seqc_S1R1r1 | 0.9900 | 0 | 0.8814 | 0 | 0.0025 | 0.0882 |
| seqc_S1R1r2 | 0.9896 | 0 | 0.8808 | 0 | 0.0025 | 0.0935 |
| seqc_S1R1r3 | 0.9913 | 0 | 0.8809 | 0 | 0.0028 | 0.0388 |
| seqc_S1R1r4 | 0.9894 | 0 | 0.8797 | 0 | 0.0024 | 0.0944 |
| seqc_S1R1r5 | 0.9901 | 0 | 0.8800 | 0 | 0.0026 | 0.0608 |
| seqc_S1R1r6 | 0.9910 | 0 | 0.8787 | 0 | 0.0027 | 0.0434 |
| seqc_S1R1r7 | 0.9890 | 0 | 0.8800 | 0 | 0.0026 | 0.0650 |
| seqc_S1R1r8 | 0.9904 | 0 | 0.8795 | 0 | 0.0023 | 0.1275 |
| seqc_S1R2r1 | 0.9839 | 0 | 0.8804 | 0 | 0.0028 | 0.0382 |
| seqc_S1R2r2 | 0.9913 | 0 | 0.8813 | 0 | 0.0025 | 0.0922 |
| seqc_S1R2r3 | 0.9877 | 0 | 0.8770 | 0 | 0.0023 | 0.1342 |
| seqc_S1R2r4 | 0.9903 | 0 | 0.8794 | 0 | 0.0026 | 0.0664 |

| Sample | Pearson | | Spearman | | Kolmorogorov-Smirnov | |
|---|---|---|---|---|---|---|
| | Statistic | P-value | Statistic | P-value | Statistic | P-value |
| seqc_S1R2r5 | 0.9917 | 0 | 0.8799 | 0 | 0.0026 | 0.0685 |
| seqc_S1R2r6 | 0.9900 | 0 | 0.8796 | 0 | 0.0027 | 0.0448 |
| seqc_S1R2r7 | 0.9889 | 0 | 0.8792 | 0 | 0.0025 | 0.0864 |
| seqc_S1R2r8 | 0.9863 | 0 | 0.8785 | 0 | 0.0027 | 0.0530 |
| seqc_S2R1r1 | 0.9928 | 0 | 0.8804 | 0 | 0.0022 | 0.1673 |
| seqc_S2R1r2 | 0.9917 | 0 | 0.8792 | 0 | 0.0025 | 0.0736 |
| seqc_S2R1r3 | 0.9916 | 0 | 0.8813 | 0 | 0.0025 | 0.0806 |
| seqc_S2R1r4 | 0.9919 | 0 | 0.8783 | 0 | 0.0028 | 0.0413 |
| seqc_S2R1r5 | 0.9933 | 0 | 0.8798 | 0 | 0.0028 | 0.0422 |
| seqc_S2R1r6 | 0.9924 | 0 | 0.8813 | 0 | 0.0024 | 0.1081 |
| seqc_S2R1r7 | 0.9920 | 0 | 0.8815 | 0 | 0.0025 | 0.0728 |
| seqc_S2R1r8 | 0.9924 | 0 | 0.8790 | 0 | 0.0025 | 0.0895 |
| seqc_S2R2r1 | 0.9921 | 0 | 0.8789 | 0 | 0.0024 | 0.1189 |
| seqc_S2R2r2 | 0.9932 | 0 | 0.8804 | 0 | 0.0023 | 0.1477 |
| seqc_S2R2r3 | 0.9924 | 0 | 0.8800 | 0 | 0.0022 | 0.1802 |
| seqc_S2R2r4 | 0.9932 | 0 | 0.8801 | 0 | 0.0025 | 0.0913 |
| seqc_S2R2r5 | 0.9914 | 0 | 0.8801 | 0 | 0.0023 | 0.1373 |
| seqc_S2R2r6 | 0.9928 | 0 | 0.8797 | 0 | 0.0026 | 0.0565 |
| seqc_S2R2r7 | 0.9913 | 0 | 0.8807 | 0 | 0.0026 | 0.0589 |
| seqc_S2R2r8 | 0.9931 | 0 | 0.8811 | 0 | 0.0026 | 0.0608 |
| **Min** | 0.9839 | 0 | 0.8770 | 0 | 0.0022 | 0.0382 |
| **Max** | 0.9933 | 0 | 0.8815 | 0 | 0.0028 | 0.1802 |
| **Mean** | 0.9909 | 0 | 0.8799 | 0 | 0.0025 | 0.0851 |
| **Median** | 0.9914 | 0 | 0.8800 | 0 | 0.0025 | 0.0771 |

### Benchmarking Results

#### Per-Rule Performance Metrics

Performance metrics were collected using Snakemake's benchmarking functionality. Values represent the mean across all samples processed in our validation analysis for three replicates. Benchmarking was conducted on a workstation with 24 cores (approximate clock speed 3 GHz) and 256 GB of RAM. The alignment was performed against the GRCh38 reference genome (Ensembl release 115).

### Supplementary Figures

#### Figure S1. AutoRNAseq Workflow Architecture

Directed acyclic graph showing dependencies between Snakemake rules. The colors surrounding each rule name is not indicative nor deterministic of any component in the workflow (e.g., intermediate files, outputs, etc.).

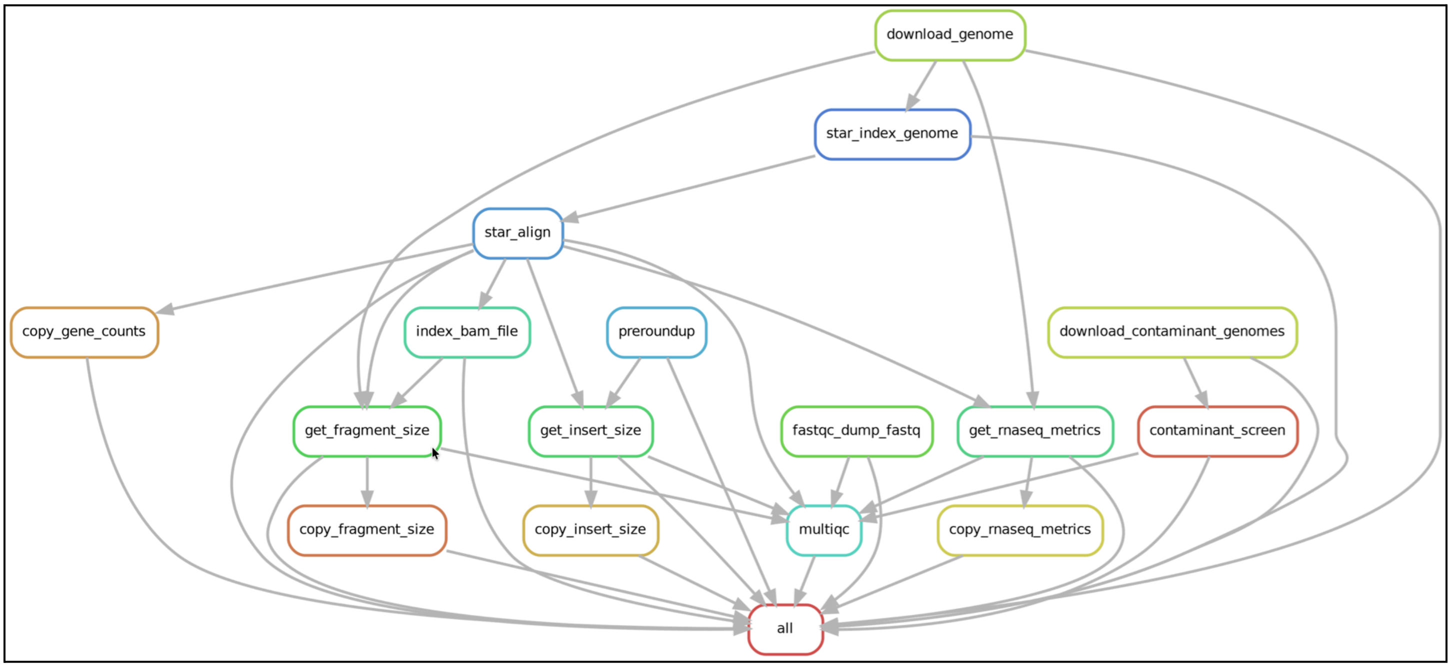
